# Inhibition of somatostatin enhances the long-term metabolic outcomes of sleeve gastrectomy in mice

**DOI:** 10.1101/2023.04.11.536368

**Authors:** Doron Kleiman, Yhara Arad, Shira Azulai, Aaron Baker, Michael Bergel, Amit Elad, Liron Hefetz, Hadar Israeli, Mika Littor, Anna Permyakova, Itia Samuel, Joseph Tam, Rachel Ben-Haroush Schyr, Danny Ben-Zvi

## Abstract

Bariatric surgery is an effective obesity treatment, leading to weight loss and improvement in glycemia, that is characterized by hypersecretion of gastrointestinal hormones. However, weight regain and relapse of hyperglycemia are not uncommon. Here, we investigated the role of somatostatin (Sst) in bariatric surgery outcomes using a mouse model of sleeve gastrectomy (SG). Sst knockout (sst-ko) mice fed with a calorie-rich diet gained weight normally, and had a mild favorable metabolic phenotype compared to heterozygous sibling controls, including elevated plasma levels of Glp1. Mathematical modeling of the feedback inhibition between Sst and Glp1 showed that Sst exerts its maximal effect on Glp1 under conditions of high hormonal stimulation, such as following SG. Obese sst-ko mice that underwent SG had higher levels of Glp1 compared with heterozygous SG-operated controls. Accordingly, SG-sst-ko mice regained less weight than controls and maintained lower glycemia months after surgery. Obese wild-type mice that underwent SG and were treated daily with a Sst receptor inhibitor for two months, had higher Glp1 levels, regained less weight, and improved glycemia compared to saline-treated SG-operated controls. Our results suggest that Sst signaling inhibition enhances and maintains the long-term favorable metabolic outcomes of bariatric surgery.

## Introduction

Somatostatin (Sst) is a neuropeptide and hormone expressed in numerous endocrine and neural cells that signals through five receptors. Sst is a negative regulator of neural activity, of both exocrine and endocrine secretion, and can inhibit cell proliferation in some cases(1). Sst secreted from the hypothalamus suppresses growth hormone (GH) secretion from the anterior pituitary gland(2–4). Sst secreted by delta cells in the endocrine pancreas inhibits insulin and glucagon secretion through paracrine and endocrine signaling(5, 6). Similarly, Sst secreted by delta cells in the stomach, intestine, and colon reduces the secretion of virtually all gastrointestinal (GI) hormones and inhibits acid secretion from gastric parietal cells(7, 8). In the clinic, Sst analogs are used to treat acromegaly, dumping syndrome, and some types of neuroendocrine tumors(9).

Despite the wide distribution of Sst-secreting cells and their effects on many organs and systems, Sst knockout mice (sst-ko) have a relatively mild phenotype. sst-ko mice display different GH secretion patterns and higher levels of GH but are not larger than wild-type mice(10). Rather, male sst-ko mice have a feminized hepatic gene expression(11, 12). sst-ko mice are hypergastrinemic, and their stomach has more parietal cells, secretes more acid, and has more proliferating cells(13). Behavioral studies demonstrated impairment in motor learning in sst-ko mice(14)

Sst deficiency is associated with lower glucose levels, especially in neonate mice (6, 15, 16). *In vitro*, islets of adult sst-ko mice have a higher basal and glucose-stimulated insulin secretion rate and secrete glucagon in response to high glucose(17). Adult sst-ko mice were reported to have lower glycemia but do not display hyperinsulinemia (4). Inhibition of Sst Receptor 2 or Sst Receptor 5, expressed in the murine intestinal L-cells, improved glucose tolerance due to Glp1 hypersecretion(18).

Sleeve gastrectomy (SG) is a common bariatric surgery wherein most of the stomach is excised along the greater curvature to produce a sleeve-shaped stomach. This surgery leads to weight loss and improved glycemic control, and alleviation of metabolic disorders in most patients and rodent models(19–23). SG affects the microbiome, the levels of bile acids and other metabolites in the blood, and leads to a post-prandial increase in GI and pancreatic hormones such as Glp1 and insulin and modulates processes that regulate systemic metabolism(24–27). SG affects gastric emptying and intestinal motility (28, 29), and in some cases causes dumping syndrome (30, 31), which can be treated by Sst analogs(32, 33). The effects of SG may wane over time, and weight regain and relapse of hyperglycemia and other metabolic diseases are not uncommon(34, 35).

In this study, we aimed to test the role of Sst in the context of obesity and SG. We found that obese sst-ko mice had a superior long-term response to SG compared to their heterozygous siblings. Pharmacological inhibition of Sst Receptors in wild-type obese mice enhanced the long-term metabolic outcomes of SG. These improvements were associated with higher levels of Glp1 in the plasma.

## Results

### Sst knockout mice gain weight normally but display better glycemic control following a high-fat high-sucrose diet

Male sst-ko mice and their heterozygous male siblings were fed a high-fat high-sucrose (HFHS) diet for 110 days. Genotype did not affect weight gain, and both groups became obese (Figure 1A, S1A). 11 weeks after starting the diet, the total energy expenditure of sst-ko mice was higher (Figure 1B). By the end of the experiment, sst-ko mice had higher fat mass and lower lean mass than littermate controls (Figure 1C-D) - with no difference in weight. There was no difference in plasma triglycerides or cholesterol in sst-ko mice and their siblings (Figure S1B-C). The livers of sst-ko mice had a lower degree of hepatic steatosis, higher glycogen content, and higher hepatic expression of genes characterizing female mice (Figure 1E-H, S1D). These results are overall consistent with previous studies characterizing sst-ko mice (10–12).

**Figure 1.**
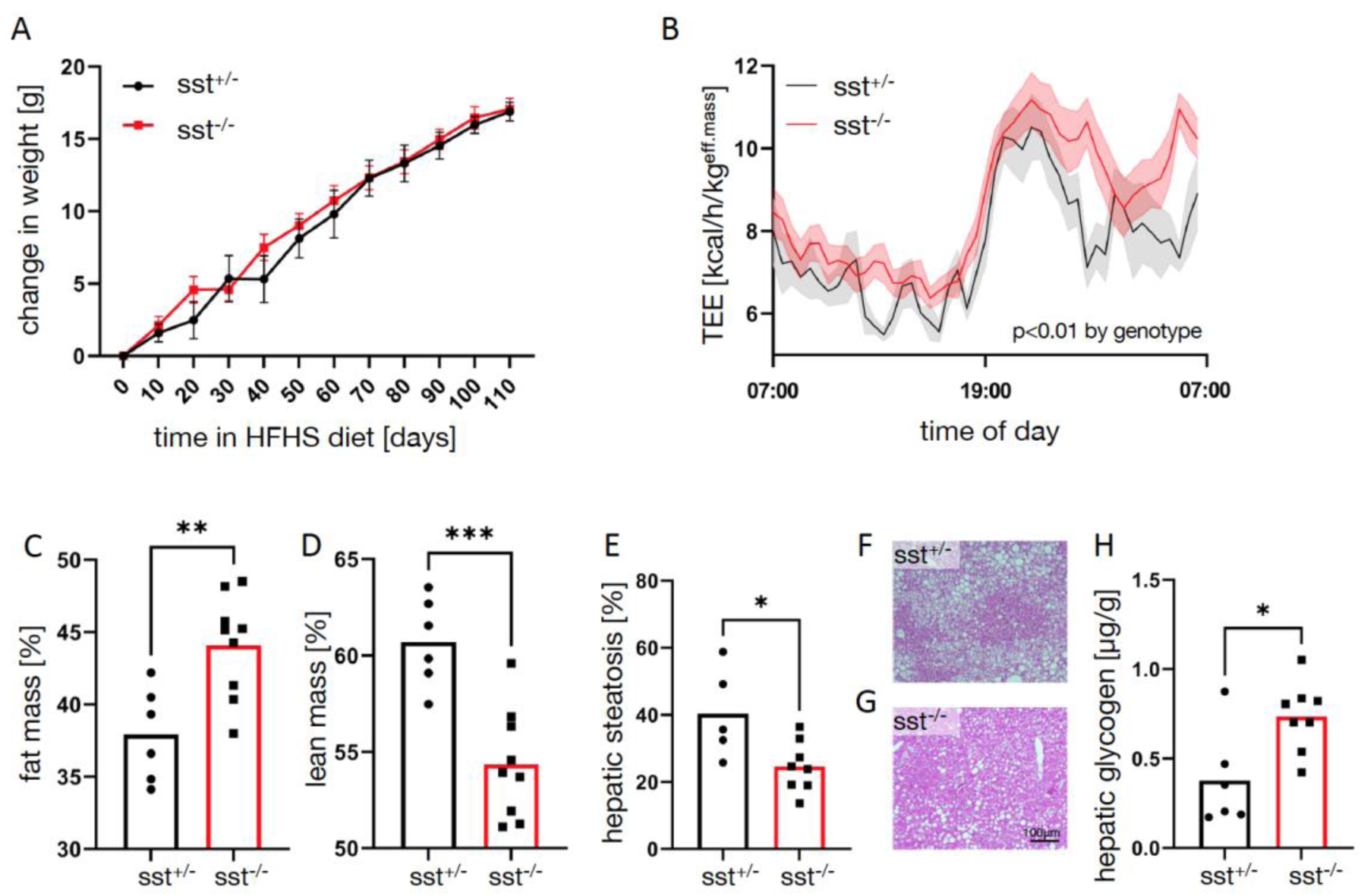
Metabolic characterization of sst-ko mice following a calorie rich diet. A. Weight gain in male sst-ko mice (red) and their male heterozygous siblings (black) during high-fat high-sucrose feeding. n=6,9. B. Total energy expenditure over 24 hours in sst-ko mice and heterozygous siblings. n=8,6. p<0.01 for genotype in 2-way continuous measurement ANOVA. C-D. Fat (C) and lean mass (D) of sst-ko mice and heterozygous siblings. E-G. Percent of lipid area in hepatic H&E stains of sst-ko and heterozygous siblings (E). A representative hematoxylin and eosin image of a liver of an sst-ko mouse is shown in (F) and of a heterozygous mouse in (G). Scale bar=100μm for F,G. H. Glycogen content in the livers of sst-ko and heterozygous siblings. *,** p<0.05, p<0.01 using Student’s t-test. Error bars in A,B denote SEM.

sst-ko mice had lower fasting and average glucose levels, and lower fasting insulin levels (Figure 2A-C). sst-ko mice also responded better to an oral glucose tolerance test (OGTT) (Figure 2D-E). To gain better insight into the glycemia of sst-ko mice, we implanted a continuous glucose measurement device on sst-ko mice and sibling controls fed on an HFHS diet and followed their glycemia for a week(36). Data showed a reduction in mean glucose levels and higher variability of glucose levels in sst-ko mice (Figure 2F-H). Mild hypoglycemia, defined as non-fasting glucose<85mg/dL was observed only in sst-ko mice. Fasting Glp1 levels increased in the plasma of sst-ko mice compared with heterozygous siblings (Figure 2I).

**Figure 2.**
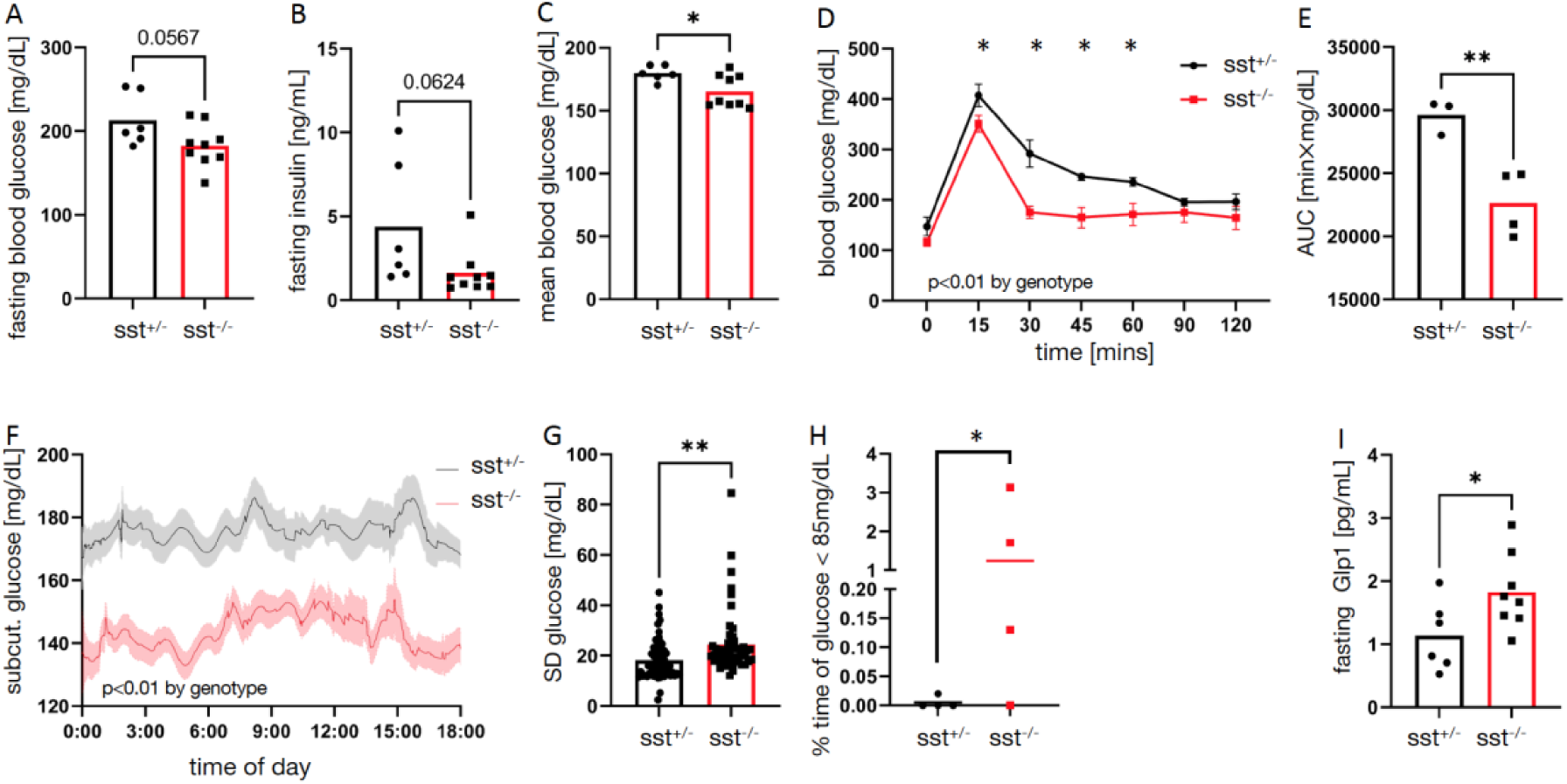
Glycemia of sst-ko mice following a calorie rich diet. A. Fasting blood glucose levels of sst-ko mice (red) and their heterozygous siblings (black). B. Fasting plasma insulin levels of sst-ko mice and their heterozygous siblings. C. Average non-fasting plasma glucose levels of sst-ko mice and their heterozygous siblings. D-E. Plasma glucose levels (D) and area under the curve (E) of sst-ko mice and their heterozygous siblings following an oral glucose tolerance test. F. Average glucose levels measured using a continuous glucose monitor device. Shades show the standard error of the mean. n=4,4 G-H. Standard deviation in glucose levels measured by a continuous glucose monitor device. Each dot represents one day in one mouse. p<0.01 by genotype in a 2-way repeated measurement ANOVA. H. The percentage of time in which glucose was lower than 85mg/dL was measured by a continuous glucose monitor device. I. Fasting plasma Glp1 levels in sst-ko mice and their heterozygous siblings. *,** p<0.05, p<0.01 using Student’s t-test. Error bars in D,F denote SEM.

### A mathematical model for the regulation of Glp1 secretion by Sst

Glp1 and Sst form a negative feedback loop wherein Glp1 increases Sst secretion, and Sst inhibits Glp1 and Sst secretion by direct interaction (Figure 3A)(18, 37, 38). The factors that induce intestinal Glp1 and intestinal Sst secretion are not entirely known. Previous studies and single-cell gene expression atlases have shown that gastrointestinal L and delta cells express many types of receptors and can metabolize luminal contents to control hormone secretion(39–43). We modeled the Glp1-Sst feedback to gain further insight into their interaction. Analytical approximations and simulations show that the effect of Sst is strongest in terms of absolute difference in Glp1 secretion and in fold-change in Glp1 secretion cases of high stimulation of Glp1 and Sst secretion (Figure 3B, S2A-B, SI text).

**Figure 3.**
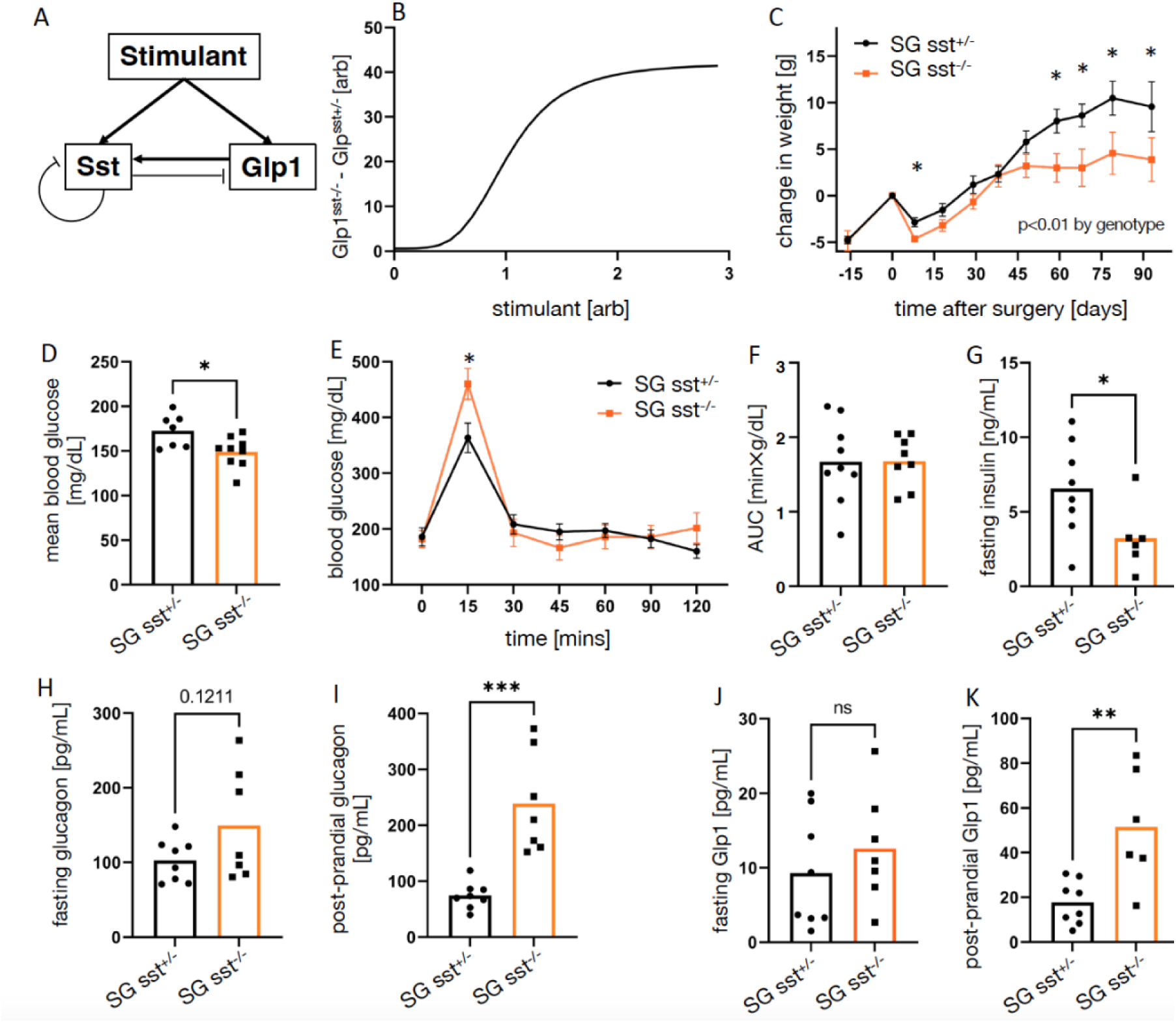
Long-term response of obese sst-ko mice to sleeve gastrectomy. A. A diagram describing the interaction between a stimulant, Glp1 and Sst. B. The difference in Glp1 level in sst-ko mice and wild-type mice as a function of the hormonal stimulation according to the mathematical model. Glp1 and stimulant levels in arbitrary units. C. Change in weight of male SG-sst-ko mice (SG-sst^-/-^, orange), and male SG-het siblings (SG-sst^+/^-, black) following surgery. n=7,9. D. Average glucose levels of SG-sst-ko mice and SG-het siblings. Each dot represents the average glucose levels of a single mouse after surgery. E-F. Blood glucose levels (E) and area under the curve (F) of sst-ko mice and their heterozygous siblings during an oral glucose tolerance test performed 6 weeks after surgery. G. Fasting plasma insulin levels. H-I. Fasting (H) and 15 minutes postprandial (I) plasma glucagon levels. J-K. Fasting (J) and 15 minutes postprandial (K) plasma Glp-1 levels. *,** p<0.05, p<0.01 using Student’s t-test. Error bars in C,E denote SEM.

### Sst-ko mice have superior metabolic outcomes following sleeve gastrectomy

SG was shown by us and many other groups to increase the secretion of Glp1 as well as other anorexigenic gut hormones in rodents(25, 26, 44). Based on our model, we hypothesized that SG in sst-ko mice will lead to an even greater secretion of Glp1, and affect the overall outcomes of surgery.

To test this hypothesis, male sst-ko mice and heterozygous littermate controls were fed an HFHS diet and underwent SG. Mice were kept on the HFHS diet for 90 more days. There was no difference in post-surgical mortality between the strains. 1-2 months after surgery, SG-sst-ko mice regained less weight than SG-het controls (Figure 3C, S2C). SG- sst-ko mice had lower average blood glucose, with no difference in fasting blood glucose (Figure 3D, S2D). Both sst-ko and controls had low-grade hepatic steatosis after SG despite the prolonged HFHS diet and a similar plasma lipid profile (Figure S3E-G). SG- sst-ko mice had higher blood glucose levels 15 minutes after oral glucose gavage but had the same glucose tolerance as SG-het controls (Figure 3E-F). Hormonally, SG-sst-ko mice had lower fasting insulin and higher post-prandial glucagon levels (Figure 3G-I). SG-sst- ko had 3-fold higher post-prandial levels of Glp1, with a mean difference of nearly 40pg/mL (Figure 3J-K).

### Pharmacological inhibition of somatostatin receptors improves metabolic outcomes of sleeve gastrectomy

The improved metabolic outcomes of SG in sst-ko mice could be a result of the different physiology of these mice before surgery which makes surgery more effective, a result of a modified response to SG, or both. To isolate the role of Sst after surgery, we used a pharmacological inhibitor of Sst signaling after SG. C57Bl6 male mice were fed an HFHS for 20 weeks. All mice underwent SG at week 11. After 1 week, the mice were randomized into a group that received daily subcutaneous injections of CycloSst (cSst), a pan-Sst receptor inhibitor, or saline(45). cSst-treated mice had a lesser weight regain and weighed less than saline-treated mice 9 weeks after surgery (Figure 4A, S4A). There was no difference in hepatic steatosis, which was very low in both groups as a result of SG (Figure S3B). cSst-treated mice did not display a reduction in a mixed-meal tolerance test performed 4 weeks after surgery but had lower fasting and post-prandial glucose levels at the end of the experiment, 9 weeks after surgery (Figure 4B-C, S3C-D). At that time, cSst mice displayed lower plasma lipid levels (Figure 4D, S3E). There was no difference in the fasting plasma insulin, c-peptide, or glucagon levels between cSst- and saline-treated SG- operated mice, yet fasting and post-prandial Glp1 levels were higher in cSst-treated mice (Figure 4E-F, S3F-H).

**Figure 4.**
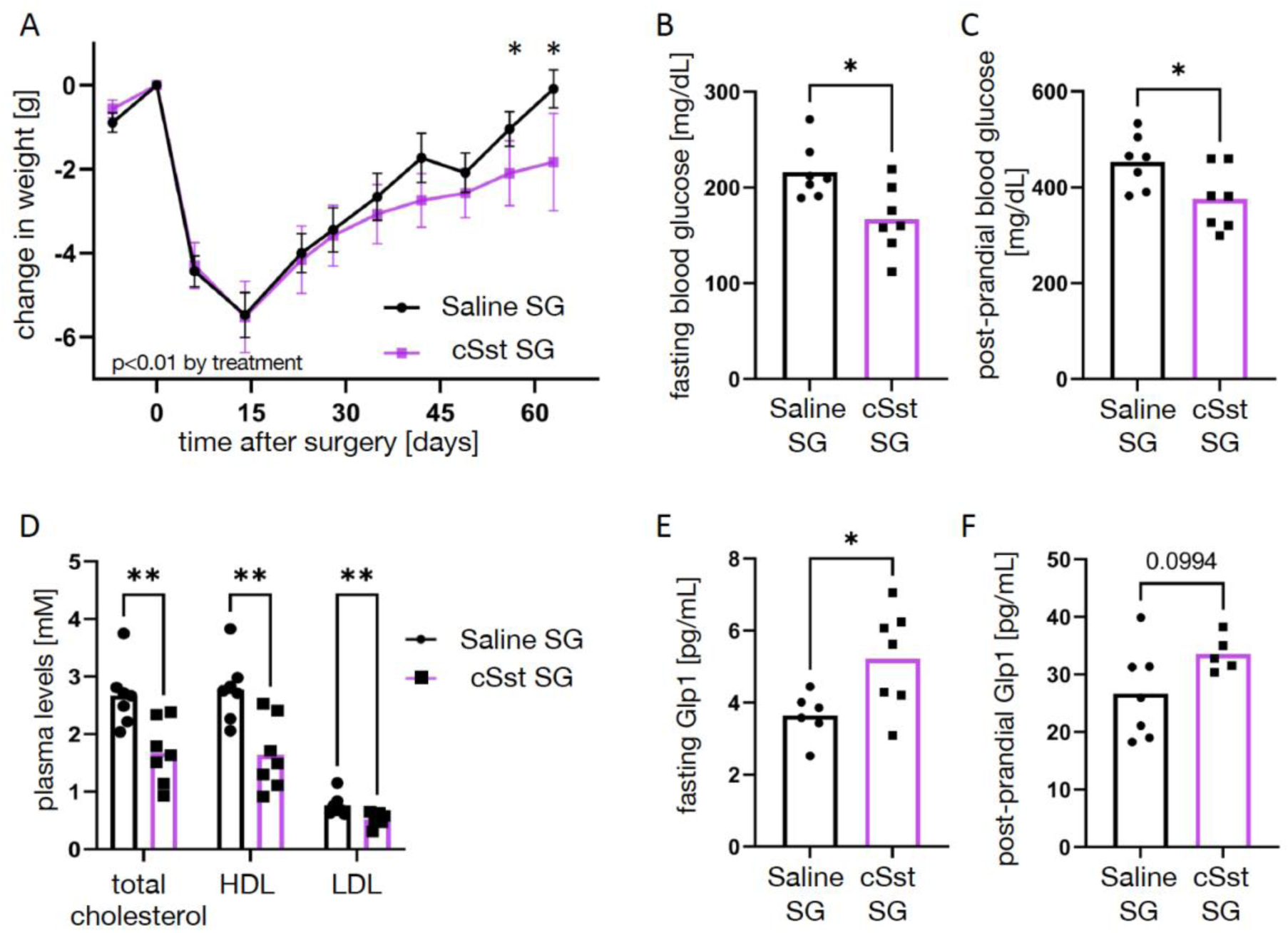
Long term response of obese mice to SG followed by inhibition of somatostatin signaling. A. Change in weight of SG-cSst mice (purple) and SG-saline mice (black). n=7,7 B-C. Fasting (B) and 15 minutes post-prandial (C) plasma glucose levels at the end of the experiment. D. Plasma total plasma cholesterol, HDL-cholesterol, and LDL-cholesterol. E-F. Fasting (E) and postprandial (F) plasma Glp-1 levels. *,** p<0.05, p<0.01 using Student’s t-test. Error bars in A denote SEM.

## Discussion

Given the wide distribution of Sst and its receptors in neurons, endocrine and exocrine cells, the phenotype of sst-ko mice is surprisingly mild(10). Using mathematical modeling we hypothesized that sst-ko mice will present a stronger phenotype in conditions where Sst-regulated hormones receive strong secretion stimuli. SG leads to the hypersecretion of many GI and pancreatic hormones that are regulated by Sst(24, 25, 27, 37). Indeed, sst-ko mice and mice treated with a Sst-receptor inhibitor displayed higher post-prandial Glp1 and glucagon levels, lower fasting insulin levels, better glycemia, and lower weight regain months after SG.

### sst-ko mice have a mild metabolic phenotype

sst-ko mice gained weight at the same rate as their heterozygous siblings, but mass distribution was different: their fat mass was higher, and their lean mass was lower. At the same time, sst-ko mice had lower-grade hepatic steatosis and higher hepatic glycogen content, suggestive of greater utilization of hepatic fatty acids, or higher lipid export from the liver to adipose tissue. These observations can be explained in part by the reported differences in GH signaling in sst-ko mice, which affect lipid distribution, hepatic metabolism, and lean mass(7, 10, 11).

We characterized the differences in glycemia between sst-ko mice and heterozygous siblings using a CGM device. sst-ko mice had lower average glycemia and a wider distribution of glucose levels. These observations are in line with the role of Sst in repressing insulin and glucagon secretion and limiting hyperglycemia and hypoglycemia(16). We noted that some sst-ko mice had non-fasting glycemia of less than 85mg/dL after a prolonged HFHS diet over 1% of the time, while heterozygous siblings did not reach these low glucose levels. The lower fasting insulin levels in sst-ko mice are counter-intuitive but can be a result of increased hepatic insulin clearance in obese sst-ko mice, and lower glycemia which reduces insulin secretion. sst-ko mice displayed an increase in fasting Glp1 levels, consistent with a tonic inhibition of Glp1 by Sst(38).

### sst-ko mice have a better long-term response to sleeve gastrectomy

Glp1 and Sst form a negative feedback loop wherein Glp1 induces or enhances Sst secretion, while Sst inhibits Glp1 secretion(18, 38). A similar feedback topology exists between insulin and Sst, although in this case, beta cells secrete Ucn3 to enhance Sst secretion. In humans, Ucn3 is also expressed in alpha cells, suggesting a similar topology in alpha cells(46). Gastrin and Sst may share a similar negative feedback topology(47). Mathematical modeling showed that in conditions of high stimulation of hormone secretion, this negative feedback would have a more pronounced effect. Based on published data, we assumed that luminal signals induce both Glp1 and Sst secretion and that Glp1 amplified Sst secretion. Assuming that Glp1 induced Sst secretion directly, or that both Glp1 and luminal contents induce Sst secretion does not affect the conclusion that Sst has the strongest inhibitory effect on Glp1 secretion in conditions of a high Glp1 stimulatory signal (SI text).

SG leads to hypersecretion of insulin, Glp1, and other GI hormones that are regulated by Sst and is therefore a unique experimental model to study enteroendocrine hormone secretion. The average fasting Glp1 level was higher in SG-operated mice compared with non-operated mice in both genotypes. Moreover, SG-sst-ko mice had 3-fold higher post-prandial Glp1 levels than SG-heterozygous mice. Over time, an increase in Glp1 and possibly other anorexigenic GI hormones can lead to lower weight and improved glycemia. Indeed, SG-sst-ko mice gained less weight and had lower glucose levels than SG- operated heterozygous controls.

SG increases glucagon levels, particularly in rodent models(20, 48). Post-prandial glucagon was elevated in SG-operated sst-ko mice compared with heterozygous controls and also compared with fasting glucagon levels in SG-sst-ko mice. High glucose stimulates glucagon secretion from islets isolated from sst-ko mice, implying that Sst secreted from pancreatic delta cells represses glucagon secretion in response to high glucose(17). The high post-prandial levels of glucagon in SG-sst-ko mice may reflect a failure of glucagon secretion inhibition in high glucose levels in this genotype. Fasting insulin levels were lower in SG-sst-ko mice, but this result can be attributed to the lower weight and glycemia of these mice. We did not measure post-prandial insulin levels due to technical reasons, yet we hypothesize that they would be high, because of the high Glp1 and glucose levels observed 15 minutes after the gavage of glucose.

### cSst-treated mice show better long-term response to sleeve gastrectomy

Alternative explanations for the superior metabolic parameters of SG-sst-ko mice are the prior lower insulin and glucose levels and different fat distribution of obese sst-ko mice compared with heterozygous siblings, or effects of neurostatin, a peptide derived from Prosomatostatin. We used cSst(45), a pan-Sst receptor inhibitor in C57Bl6 mice to control for pre-existing differences between sst-ko mice and their heterozygous siblings. By randomizing the mice into cSst- or saline-treated mice a week after SG, we also controlled for differences in the early response to surgery.

cSst-SG mice maintained higher weight loss and had better glycemia than saline-SG controls, supporting the results from the genetic model. They had similar levels of insulin and glucagon, and similar low-grade hepatic steatosis, yet displayed higher levels of plasma Glp1. These results support a role for post-surgical Sst signaling in regulating the long-term outcomes of SG. This preliminary study suggests that Sst inhibitors may enhance the long-term outcomes of SG and possibly other bariatric surgeries in pre- clinical settings. Importantly, we studied here male mice only and did not include a post-surgical dietary intervention.

### Mechanisms for maintenance of weight loss and low glycemia after SG

Given the significance of Glp1 in the physiology of bariatric surgery, we hypothesize that the outcomes of SG in sst-ko or cSst-treated mice can be attributed in part to Glp1. However, Sst inhibits the secretion of virtually all GI and pancreatic hormones, as well as GH. Bariatric surgery is associated with hypersecretion of GI and pancreatic hormones, and a reduction in GH signaling(27, 49, 50). Moreover, SG was shown to have the same metabolic effects in wild-type and glp1r-ko mice, demonstrating the role of other factors in mediating the outcomes of surgery(51). Improvement in the long-term metabolic outcomes of SG by Sst inhibition or deletion is likely not only a result of an increase in Glp1 signaling.

The full knockout genetic model and the pan-receptor inhibitor we used do not allow us to determine which hormone or what endocrine, exocrine, or neural system is most important for enhancing the metabolic outcomes of SG. Since insulin and glucagon levels were not affected by cSst treatment we believe that the effect of cSst on the endocrine pancreas is not a major determinant of the surgical outcome. We also note that, unlike SG-sst-ko mice, cSst-treated SG-operated mice did not display high post-prandial glucose levels, suggesting that a dumping syndrome was not induced following cSst treatment. Glp1 levels however were affected by cSst treatment, and we hypothesize that alleviation of Glp1 inhibition by Sst may underlie the phenotype we observed after SG.

The effects of sst-ko and cSst treatment become apparent 1-2 months after surgery. One explanation is that the long-term effect relates to a Sst-mediated adaptive process that occurs normally after SG, such as changes in GH signaling (25, 49, 50). It is also possible, that the short-term effects of surgery are not highly dependent on Glp1 or other hormones regulated by Sst, but are more affected by mechanical and entero-neuronal effects of surgery, while the long-term effects are more sensitive to metabolic, hormonal, and neuronal signaling that are regulated by Sst(26). The simplest explanation is that the increase in Glp1 or other Sst-regulated hormones generates a small effect with a detectable systemic phenotype only months after surgery.

## Conclusion

Bariatric surgery is an aggressive yet effective treatment for obesity and metabolic diseases. The long-term outcomes of surgery vary between patients: while many patients display significant long-lasting weight loss, some patients experience partial or even full weight regain years after surgery(21). Patients that regain weight or remain obese after bariatric surgery can be treated by lifestyle modifications, behavioral therapy, Glp1 receptor agonists and other anti-obesity treatments(34, 35). Here, we provided evidence in male mice that genetic loss of Sst or pharmacological inhibition of Sst signaling improves the long-term metabolic outcomes of SG. Specifically, SG-operated mice with no or reduced Sst signaling had lower glycemia, lower weight, and higher Glp1 levels compared with non-treated SG-operated mice months after surgery.

## Materials and Methods

### Mice

Animal experiments were approved by the Hebrew University’s Institutional Animal Care and Use Committee (IACUC). Mice were housed in a specific pathogen-free facility on a strict 12-h light–dark cycle. For the metabolic phenotyping six-week-old *Sst^tm1Ute^* (Jackson Labs #:003117) male mice were fed on a high-fat high-sucrose (HFHS) diet (Envigo Teklad diets TD08811) for 110 days. For SG in sst-ko mice, six-week-old *Sst^tm1Ute^* male mice were fed an HFHS diet until they reached an average weight of 35 grams. Mice were then operated on and maintained over 3 months after surgery on an HFHS diet. For the pharmacologic intervention, six-week-old C57BL/6JOlaHsd male mice were fed an HFHS diet for 11 weeks, operated and maintained over 2 months after surgery on an HFHS diet.

### Surgery

Mice underwent SG surgery as described previously(19). Briefly, a midline incision through the skin and underlying linea alba was performed, followed by exposure and mobilization of the stomach. A 12 mm clip was placed horizontally across the stomach’s greater curvature using a Ligaclip Multiple Clip Applier. The excluded part of the stomach was excised, the abdominal wall was sutured using 6-0 coated vicryl sutures (Ethicon, J551G), and the skin was closed with clips (Autoclip system, FST-12020-00). Sham surgeries included the abdominal incision, exposure and mobilization of the stomach, and closure of the body wall and skin. Each surgery lasted approximately 15–20 minutes. Mice were fasted overnight the day before surgery and during the day of surgery, and then returned to the HFHS diet.

### Cyclosomatostatin treatment

Cyclosomatostatin (cSst) was purchased from Tocris (Cat. No. 3493) and dissolved in 0.8% EtOH and saline according to the manufacturer’s instructions. cSst was injected subcutaneously at 30 μg/kg body weight and injected subcutaneously every evening from 7 days after the surgery and until the end of the experiment.

### Blood glucose, oral glucose tolerance tests (OGTT) and CGM

Blood glucose was measured with a glucometer Accu-Check (Roche Inc.) by tail bleeding. Non-fasting blood glucose refers to blood glucose level at 7-8 am when animals had *ad libitum* access to food and water throughout the night. OGTT was performed on mice fasted for 6 hours between 7 am and 1 pm by oral gavage of 20% glucose solution in saline 2g/kg body weight. Blood glucose was measured at 15, 30, 45, 60, 75, 90 and 120 minutes after injection. Continuous glucose measurements (CGM) were performed using the FreeStyle Libre system by Abbot according to a recently published protocol(36). Briefly, the CGM sensor was inserted through a small incision under the skin of the back of the mouse. The CGM was attached to the skin using sutures, and glucose was measured continuously for an average of 9-10 days. The sensor stores data for up to 8 hours, and we had access to measurements between midnight and 6pm.

### Hormone measurement

Basal plasma hormones were measured following a 6-hour fast between 7 am and 1 pm. Postprandial plasma hormones were measured following a fast and 15 minutes after an oral glucose gavage as described above. Hormones were measured using Ultra-Sensitive Mouse Insulin ELISA (Crystal Chem, #90080), Mouse Glucagon ELISA (Crystal Chem, #81518), Mouse C-peptide ELISA Kit (Crystal Chem, #90050) and V-Plex Total GLP-1 kit (Mesoscale, K1503PD-2).

### Plasma lipid analysis

Plasma samples were analyzed for triglycerides, total cholesterol, LDL, HDL using a Cobas c111 (Roche Diagnostics) automated clinical chemistry analyzer that was calibrated according to manufacturer guidelines.

See SI text for additional Materials and Methods.

### Mathematical model

The following equations were used to model the interaction between Glp1 (H) and Sst (S). See SI text for details

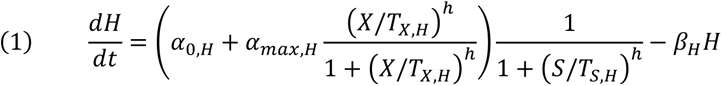

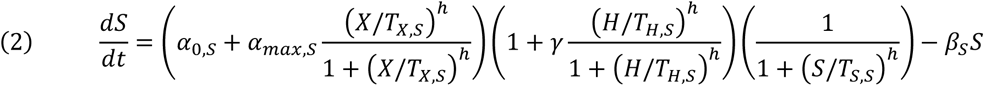

### Statistics

Student’s t-test was performed to test statistical significance when two groups were compared. Two-way repeated-measures ANOVA was performed when a time series measurement was taken. Calculations were performed in Graphpad prism9 or R-studio.

## Acknowledgments

We thank our group members for fruitful discussions. This study was funded by an ERC StG (803526) and ISF research grant (967/18) awarded to DBZ, and ISF research grant (158/18) awarded to JT. DBZ is a Zuckerman STEM faculty fellow.

## Supporting Information

### 1. Analytical analysis of the mathematical model

The following equations were used to model the negative feedback between a hormone (H) and Somatostatin (S)

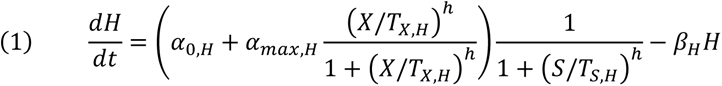

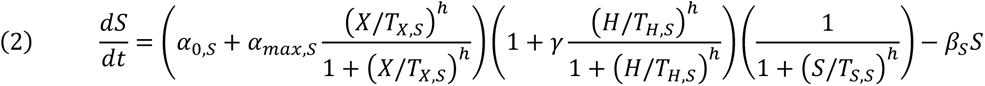

Equation (1) describes the stimulation of secretion of hormone H by a signal X, and the repression of secretion by Somatostatin. The hormone is degraded linearly. We assumed Sst represses total hormone secretion: stimulated and basal.

Equation (2) describes the stimulation of Somatostatin secretion by the same signal, enhancement of secretion by the Hormone, and auto repression. Somatostatin is degraded linearly. We use the same Hill coefficient in the equations for simplicity. Other applications of induction, enhancement and repression of secretion do not change the results qualitatively.

The model uses standard Hill function to model induction and repression.

The following parameters used in the equations and simulations. [C] is arbitrary concentration unit.

| Parameter | Interpretation | Value used in simulation |
| --- | --- | --- |
| $H$ | Levels of the hormone regulated by Sst | |
| $S$ | Somatostatin levels | |
| $\alpha_{0,H}$ | Basal hormone secretion rate | 10[C] sec<sup>-1</sup> |
| $\alpha_{max,H}$ | Maximal stimulated hormone secretion | 90 [C] sec<sup>-1</sup> |
| $X$ | Hormonal stimulation signal | Between 0-3 [C] |
| $T_{X,H}$ | Signal value in which half the maximal hormone secretion is reached | 1[C] |
| $h$ | Hill function coefficient | 4 |
| $T_{S,H}$ | Somatostatin level in which the total Hormone secretion is repressed by 50% | 4[C] |
| $\beta_H$ | Hormone degradation rate | 2sec<sup>-1</sup> |
| $\alpha_{0,S}$ | Basal Somatostatin secretion rate | 1[C] sec<sup>-1</sup> |
| $\alpha_{max,S}$ | Maximal stimulated Somatostatin secretion | 9[C] sec<sup>-1</sup> |
| $T_{X,S}$ | Signal value in which half the maximal Somatostatin secretion is reached | 1[C] |
| $\gamma$ | Factor for stimulation of Somatostatin secretion by the Hormone | 3 |
| $T_{H,S}$ | Hormone value in which stimulation of Sst secretion is 50% of its maximal value | 4[C] |
| $T_{S,S}$ | Somatostatin level in which it represses its own secretion | 4[C] |
| $\beta_S$ | Somatostatin degradation rate | 1[C] sec<sup>-1</sup> |

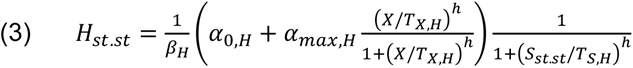

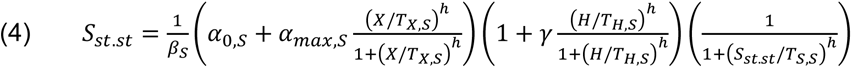

We can use the steady state assumption in both fasting and post-prandial conditions since both Glp1 and Somatostatin have a very short half-life compared to the signal (meal). Steadt state was reached in all simulations in time *t* ≫ max (*β*_*s*_^−1^, *β*_*H*_^−1^)

The steady state analysis shows that if Somatostatin levels are high enough *S*_*st*.*st*_ > *T*_*S*,*H*_, the steady state levels of the hormone would be close to zero. This may be the case for perfusion with somatostatin. However, in our simulation, we set *s*_*st*.*st*_ ≤ *T*_*s*,*H*_, so that *H*_*st*.*st*_ would be monotonous wrt *X*. This is probably not the case for *H* =Glucagon, but is the case for other hormones such as insulin and Glp1.

For the sst-ko, S=0 and (3) becomes:

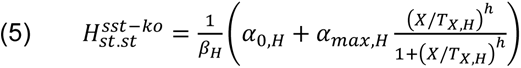

We define *ζ* as the ratio between *H*_*st*.*st*_ in sst-ko mice and in wild-type mice.

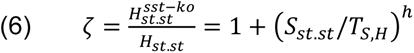

Since Sst inhibits Glp1, *ζ* ≥ 1, as expected. To understand how *X* affects *ζ*, note that

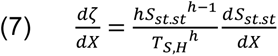

Both terms in (6) are positive: *s*_*st*.*st*_ is greater than 0, and 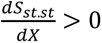 as Somatostatin levels increase with the signal according to available experimental data. The stronger the signal, the greater the fold-change effect Somatostatin has on the levels of the hormone it regulates. This result holds also if we change the functional form of S, and is independent of how the hormone affects Sst secretion.

In our data under small *X* (fasting), *ζ* ≈ 1.5, and under maximal stimulation (post-prandial in SG operated mice) *ζ* ≈ 3

The net effect of removing Sst can be quantified by

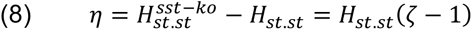

If both *H*_*st*.*st*_ and *ζ* − 1 are positive and monotonously increasing with *X*, *η* is positive and monotonously increasing in X. Therefore, maximal effect of Sst will be observed under high stimulation.

Analyzing *η* directly

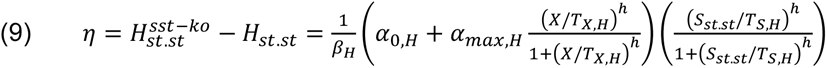

*η* takes the form of the product of two sigmoidal functions.

Basal conditions: If *X* ≪ *T*_*X*,*H*_, *T*_*X*,*S*_, and assuming low/no autorepression of Sst under low stimulation,

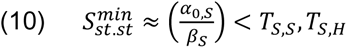

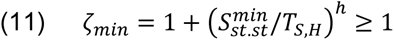

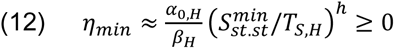

High stimulation conditions: If *X* > *T*_*X*,*H*_, *T*_*X*,*S*_, assuming high stimulation of S by H and autorepression of S:

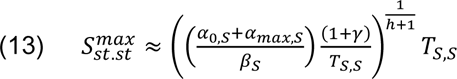

Assuming *s*_*st*.*st*_ ≈ *T*_*s*,*H*_, under maximal stimulation, we obtain

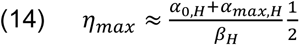

The values used in the simulation were derived using the analysis above.

using the calculated value of 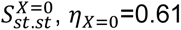 in the simulation, and 0.69 according to our analytical approximation. Similarly, *η*_*X*=3_=41.53 in the simulation, and 50 by the approximation.

Note that we did not aim to fit the many parameters of Eq. 1-2 to our data, and strived to keep the parameters as simple as possible (same threshold for secretion, same thresholds for stimulation/repression, same fold change in basal to maximal secretion, same Hill parameter).

## 2. Supporting Methods

### Terminal Blood Collection

Mice were anesthetized using ketamine 100mg/kg and xylazine 8mg/kg diluted in 0.9% sodium chloride. Blood was extracted via terminal bleeding using heparin-coated syringes and 25G needles and transferred to lithium heparin-coated tubes (Greiner, MiniCollect, #450535). Aprotinin (Sigma-Aldrich, A6279), DPP4 inhibitor (Merck #MDPP4), and EDTA (final concentration 0.83mM) were added to the blood sample to avoid degradation of glucagon and GLP-1. Blood was then centrifuged at 6,000 RCF for 1.5 minutes. Plasma was then collected and stored in liquid nitrogen until transferred to final storage at –80°C.

### RNA extraction

Total RNA was extracted from whole livers using TRI reagent (Sigma-Aldrich, T9424) according to the manufacturer’s instructions. Approximately 60 mg of tissue was used for each sample.

### qPCR

the following primers were used

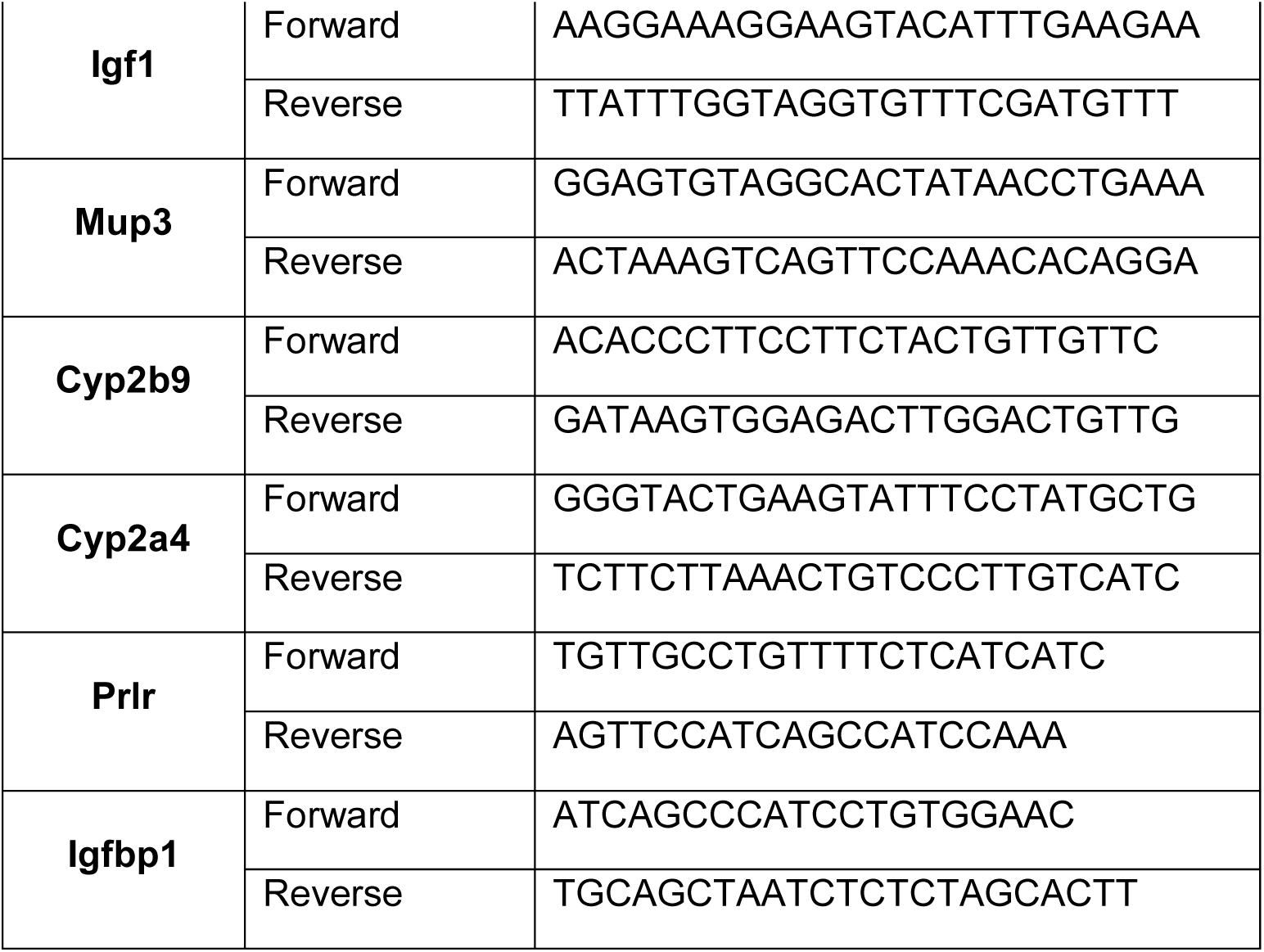

### Tissue fixation and processing

Liver samples were fixed overnight at 4°C in 4% formaldehyde, then washed in PBS and continued fixation in 70% ethanol. Later, tissue paraffinization was performed in the Hadassah Medical Center Pathology Department and then transferred to paraffin blocks. Tissues were cut into 4μm slides and deparaffinized using Xylene and decreasing percentages of alcohol.

### Tissue Staining

For the H&E stain tissues were deparaffinized and stained as previously described(20). Dried slides were sealed with Entellan mounting medium (Mercury, 1079600600). Percent of hepatic lipid droplets was calculated by a custom R script. For the Alcian blue-PAS stain paraffin slides were deparaffinized and stained with a PAS staining kit (Merck, 101646,101647) according to the manufacturer’s protocol.

### Glycogen Assay

Livers samples were extracted during the terminal procedure, frozen in liquid nitrogen and stored at –80°C. Samples were thawed on ice and glycogen content was analyzed according to the manufacturer’s instructions (Abcam, #ab65620).

### Metabolic Cages

The metabolic profiles of the mice were assessed by using the Promethion High-Definition Behavioral Phenotyping System (Sable Instruments, Inc., Las Vegas, NV, USA). Data acquisition and instrument control were performed using MetaScreen software version 2.2.18.0 (Sable Instruments, Inc., Las Vegas, NV, USA), and the obtained raw data were processed using ExpeData version 1.8.4 using an analysis script detailing all aspects of data transformation. Mice with free access to food and water were subjected to a standard 12 h light/12 h dark cycle, which consisted of a 48 h acclimation period followed by 24 h of sampling. Respiratory gasses were measured by using the GA-3 gas analyzer (Sable Systems, Inc., Las Vegas, NV, USA) using a pull-mode, negative-pressure system. Airflow was measured and controlled by FR-8 (Sable Systems, Inc., Las Vegas, NV, USA), with a set flow rate of 2000 mL/min. Water vapor was continuously measured and its dilution effect on O2 and CO2 was mathematically compensated. Respiratory quotient (RQ) was calculated as the ratio between CO2 produced (VCO2) to O2 consumed (VO2) based on Equation:RQ = VCO2/VO2. Total energy expenditure (TEE) was calculated using VO2 and RQ based on the Equation: TEE = VO2 × (3.815 + 1.232 × RQ).

## 3. Supporting Figures

**Figure S1.**
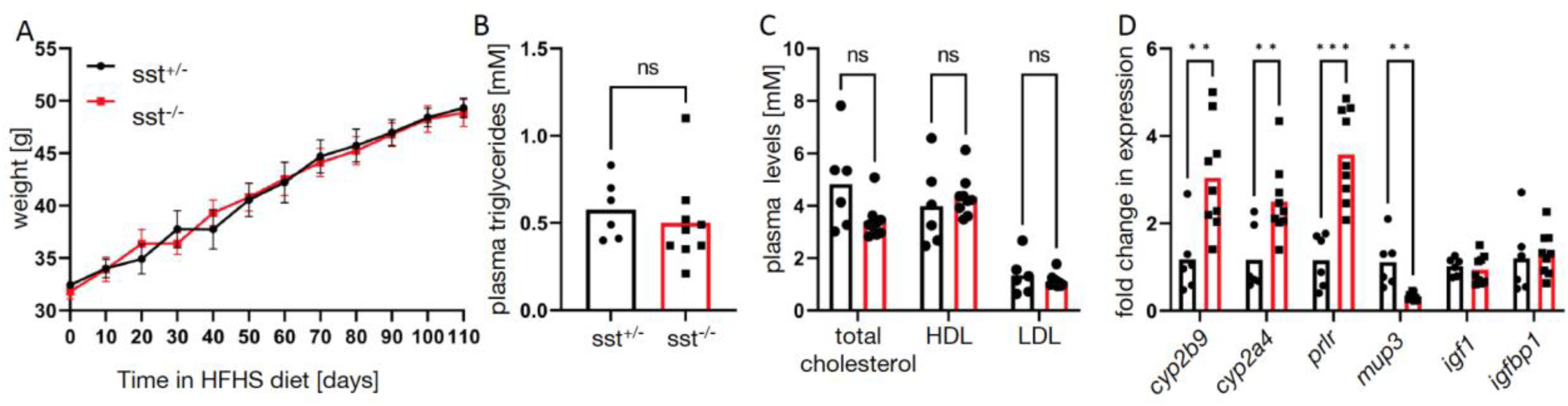
supporting Figures 1,2. A. Weight of male sst-ko mice (sst-/-, red) and their male heterozygous siblings (sst+/-, black) during high-fat high-sucrose feeding. n=6,9. B. Plasma triglyceride levels in sst-ko mice and heterozygous siblings. C. Plasma total cholesterol, HDL-cholesterol and LDL-cholesterol in sst-ko mice and heterozygous siblings. D. Fold change in gene expression in the livers of sst-ko mice compared with heterozygous siblings. *cyp2b9, cyp2a4, prlr* and *mup3* have higher expression in livers of female mice. There was no change in *igf1* or *igfbp1* expression. Both genes are regulated by growth hormone signaling.

**Figure S2.**
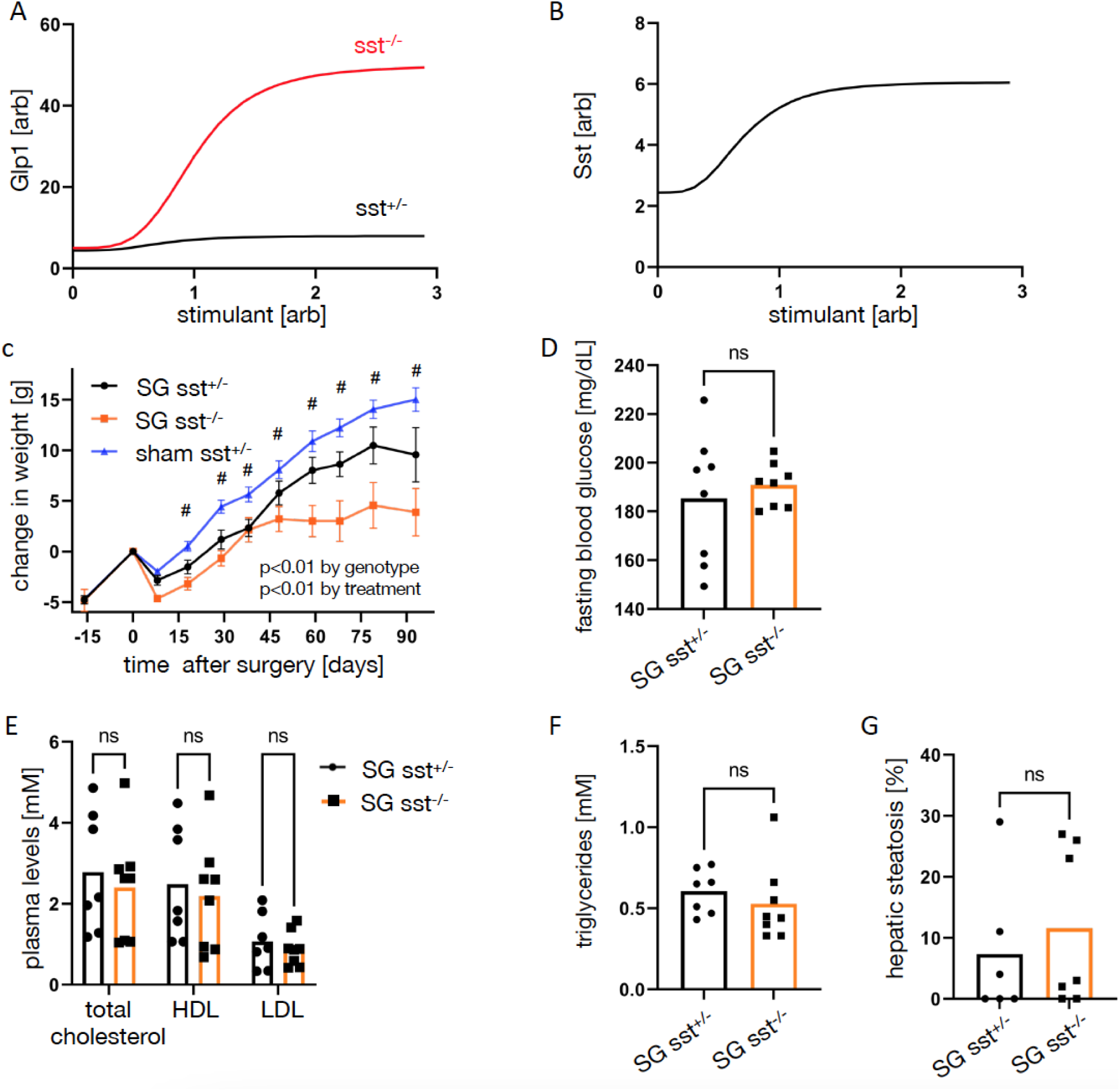
supporting Figure 3. A. Levels of Glp1 in sst-ko mice (red) and wild type mice (black) as a function of a stimulant, according to the mathematical model. B. Sst levels as a function of the stimulant according to the mathematical model. C. Change in weight of male SG-sst-ko mice (orange), male SG-het siblings (black), and sham-het siblings (blue) following surgery. n=7,9,6 D. Fasting blood glucose levels of SG-sst-ko mice and their SG-heterozygous siblings. E. Plasma total cholesterol, LDL-cholesterol, and HDL-cholesterol levels of male SG-sst- ko mice and male SG-het siblings. F. Plasma triglyceride levels of male SG-sst-ko mice and male SG-het siblings. G. Quantification of hepatic steatosis in SG-sst-ko mice and male SG-het siblings.

**Figure S3.**
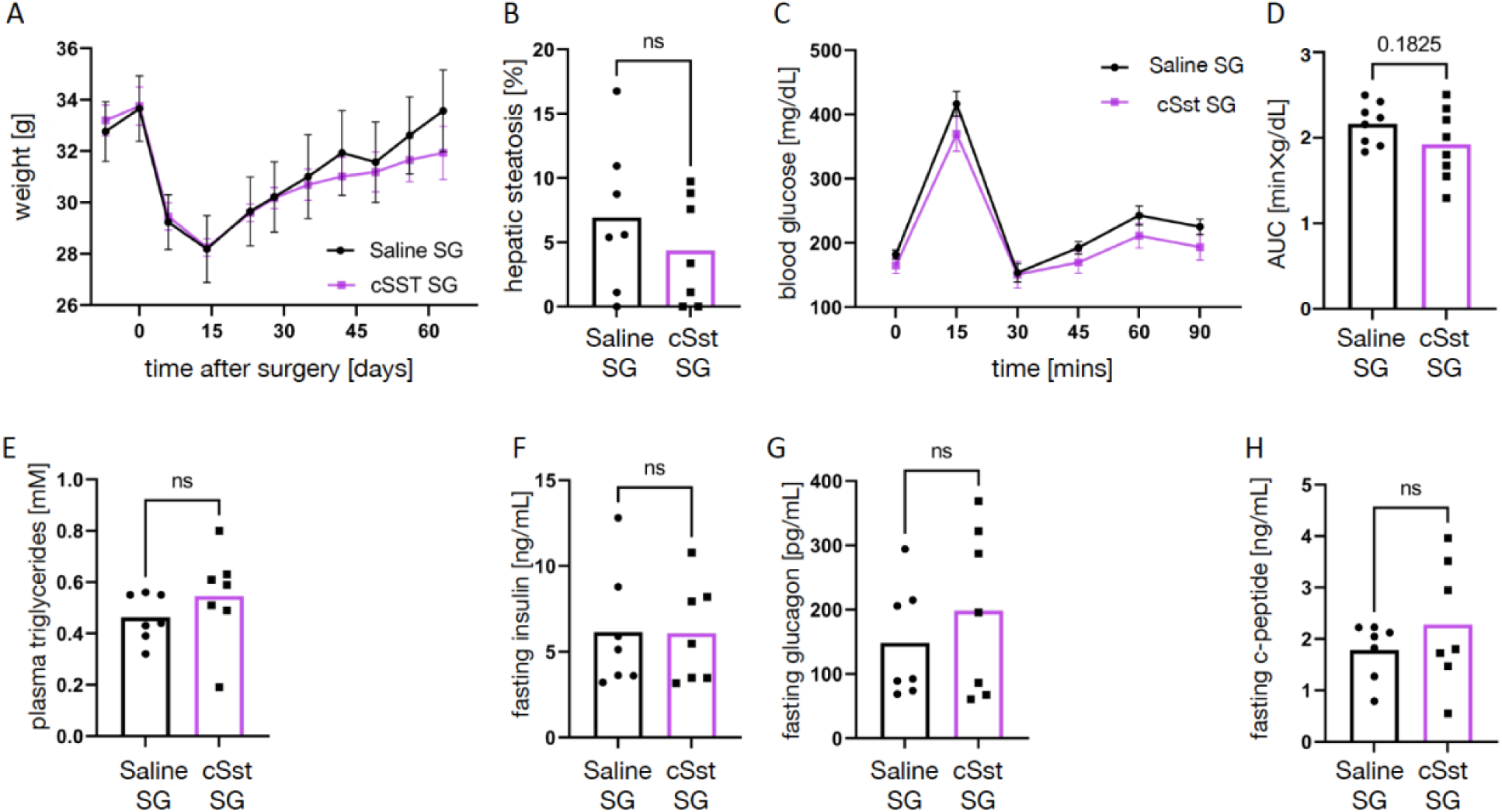
supporting Figure 4. A. Weight of SG-cSst mice (purple) and SG-saline mice (black). n=7,7. B. Percentage of the hepatic area occupied by lipids in H&E staining. C-D. Glucose levels following a mixed meal glucose tolerance test performed 4 weeks after treatment (C), and area under the curve (D). E. Fasting plasma triglycerides levels. F-H. Fasting plasma insulin (F), glucagon (G) and c-peptide (H) levels.

## Notes

**Competing Interest Statement:** The authors declare no conflict of interest.

### Competing Interest Statement

The authors have declared no competing interest.

